# Diversification of *B. subtilis* during experimental evolution on *A. thaliana* and the complementarity in root colonization of evolved subpopulations

**DOI:** 10.1101/2021.03.06.434191

**Authors:** Christopher Blake, Mathilde Nordgaard, Gergely Maróti, Ákos T. Kovács

**Affiliations:** Bacterial Interactions and Evolution Group, DTU Bioengineering, Technical University of Denmark, 2800 Kongens Lyngby, Denmark; Institute of Plant Biology, Biological Research Centre of the Hungarian Academy of Sciences, 6726 Szeged, Hungary

**Author notes:** For correspondence.; Søltofts Plads Building 221, 2800 Kgs Lyngby, Denmark.

**Keywords:** *Bacillus subtilis*, experimental evolution, root colonization, biofilm, motility, morphotype

## Abstract

The soil bacterium *Bacillus subtilis* is known to suppress pathogens as well as promote plant growth. However, in order to fully exploit the potential as natural fertilizer, we need a better understanding of the interactions between *B. subtilis* and plants. Here, *B. subtilis* was examined for root colonization through experimental evolution on *Arabidopsis thaliana*. The populations evolved rapidly, improved in root colonization and diversified into three distinct morphotypes. In order to better understand the adaptation that had taken place, single evolved isolates from the final transfer were randomly selected for further characterization, revealing changes in growth and pellicle formation in medium supplemented with plant polysaccharides. Intriguingly, certain evolved isolates showed improved root colonization only on the plant species they evolved on, but not on another plant species, namely tomato, suggesting *A. thaliana* specific adaption paths. Finally, the mix performed better than the sum of its constituents in monoculture, which was demonstrated to be caused by complementarity effects. Our results suggest, that genetic diversification occurs in an ecological relevant setting on plant roots and proves to be a stable strategy for root colonization.

**Significance Statement:** Understanding how plant-growth-promoting rhizobacteria (PGPR) colonize plant roots is crucial to fully utilize their potential for agricultural applications. Here, we employ experimental evolution of the PGPR *Bacillus subtilis* on *Arabidopsis thaliana* to study root colonization. We revealed that evolving populations rapidly improve in root colonization and diversify into distinct morphotypes. Notably, improved root colonization by evolved isolates was observed on *A. thaliana*, not on tomato. Moreover, isolates of distinct morphotypes interacted during root colonization and the mixture of morphotypes showed higher productivity than predicted. These findings suggest that genetic diversification might be a stable strategy to maximize root colonization.

## Introduction

The interplay between root-associated microbial communities and the plant itself is essential for the productivity and health of the plant (Vandenkoornhuyse *et al*., 2015). In general, these microbial communities are highly diverse and comprise various complex interactions, both beneficial as well as deleterious (Trivedi *et al*., 2020). The plant host species tends to be one of the main drivers of variation in the composition of the plant root microbiome, indicating an active selection of microbes by the plant (Micallef *et al*., 2009; Fitzpatrick *et al*., 2018). Especially beneficial microbes, e.g. plant growth promoting rhizobacteria (PGPR), are suggested to be actively recruited from the surrounding bulk soil (Yi *et al*., 2011; Pieterse *et al*., 2014).

One widely studied PGPR is the Gram-positive, non-pathogenic, spore-forming bacterium *Bacillus subtilis* (Cazorla *et al*., 2007; Todorova & Kozhuharova, 2010; Luo *et al*., 2015). *B. subtilis* is commonly found in soil and has been isolated from various plant species (Cazorla *et al*., 2007). It is widely recognized to aid plants through multiple direct as well as indirect manners (Blake *et al*., 2021). *B. subtilis* is known to promote plant growth either through production of phytohormones or by improving nutrient availability (Arkhipova *et al*., 2005; Saravanakumar *et al*., 2011; Sivasakthi *et al*., 2014). In addition, *B. subtilis* protects the plant against pathogens, either directly by competing for space and nutrients and by producing antimicrobials, or indirectly by triggering induced systemic resistance (Lugtenberg & Kamilova, 2009; Pieterse *et al*., 2014). One of the main mechanisms of plants to attract and select for *B. subtilis*, is the secretion of root exudates, which act as chemoattractant and nutrient source for microbes (Zhang *et al*., 2014). Subsequently, *B. subtilis* needs to evade the innate immune response of the plant and tightly attach to the root, often in the form of biofilms, providing further opportunities for co-evolution with their host plant (Chen *et al*., 2013; Deng *et al*., 2019). Even though some *B. subtilis* strains are already applied as biocontrol agents within agriculture, its success under field conditions varies immensely (Peng *et al*., 2011; Wei *et al*., 2016). The main cause of this variation, has been suggested to be the inability of *B. subtilis* to colonize and persist in the rhizosphere for longer periods of time (Shoda, 2000).

Up until now, most studies analyzing the importance of bacterial properties for successful root colonization by *B. subtilis* used either constructed mutant strains or distinct natural isolates (Beauregard *et al*., 2013; Chen *et al*., 2013; Allard-Massicotte *et al*., 2016; Dragoš *et al*., 2018a; Nordgaard *et al*., 2021). This revealed, amongst other things, that the capability of *B. subtilis* to colonize plant roots is tightly linked to its chemotactic response towards root exudates, as well as its ability to form biofilms (Beauregard *et al*., 2013; Chen *et al*., 2013; Allard-Massicotte *et al*., 2016). Indeed, *B. subtilis* strains exhibiting strong pellicle formation - a type of biofilm formed at the liquid-air interface – have been observed to show enhanced root colonization, while weak pellicle forming strains showed reduced root colonization (Chen *et al*., 2013). Biofilms consist of cells, that are packed tightly together, embedded in a self-produced extracellular matrix (ECM), which for *B. subtilis* consists mainly of an exopolysaccharide (EPS) and the protein TasA (Branda *et al*., 2006; Arnaouteli *et al*., 2021). Both, Beauregard *et al*. (2013) and Dragoš *et al*. (2018a) have shown that EPS and TasA are required for successful root colonization, as mutants, Δ*eps* and Δ*tasA*, displayed a reduced number of cells on plants, compared to the WT. On the contrary, inoculation of plant roots with a mixture of Δ*eps* and Δ*tasA* led to a significantly higher root colonization compared to the WT, as matrix components are shared and division of labor occurs between complementing Δ*eps* and Δ*tasA* mutants (Beauregard *et al*., 2013; Dragoš *et al*., 2018a). Matrix components such as TasA and EPS are regarded as public goods, i.e. compounds that are costly to produce but provide a collective benefit for the overall population (Branda *et al*., 2006; Zhang *et al*., 2015; Dragoš *et al*., 2018a). Public goods like matrix components, however, not only provide the basis for division of labor, where resources are used and shared efficiently, but might also allow exploitation, where cheaters are able to utilize the public good without contributing to its production (van Gestel *et al*., 2014; Dragoš *et al*., 2018a, Dragoš *et al*., 2018b).

Recently, a new study emerged using experimental evolution to analyze bacterial traits important for plant microbial interactions (Batstone *et al*., 2020). In general, experimental evolution might be less constrained by preconceptions and is a powerful tool to analyze how microorganisms adapt to a wide range of environments as well as to study evolutionary trade-offs (Kawecki *et al*., 2012). By allowing cells to grow until the population reaches a high density, and subsequently transferring only a part of the population to new, fresh medium, continued growth is granted and therefore natural selection will drive the population over generations to adapt to the given conditions (McDonald, 2019). Batstone *et al*. (2020) used experimental evolution to evolve *Ensifer meliloti* on distinct *Medicago truncatula* genotypes in a greenhouse system in soil for five plant generations. *E. meliloti* rapidly adapted to its local host genotype, meaning that derived bacteria were both more beneficial and achieved higher fitness on the plant host genotype they shared an evolutionary history with (Batstone *et al*., 2020).

In this study, we established a laboratory model system to inspect the evolution of *B. subtilis* on plant roots. 14-16d old *A. thaliana* seedlings were inoculated with *B. subtilis* in a minimal medium, after 2 days, the colonized root was subsequently transferred to fresh medium containing a new sterile seedling (14-16d old). This allowed us to follow evolving populations for 20 transfers and select for a regular root recolonization, including chemotaxis, biofilm formation and dispersal. We observed that evolving populations rapidly improved in root colonization and quickly diversified into distinct morphotypes. When these morphotypes were mixed together, they interacted on the root, and the mixed community showed “overyielding” compared to the predicted productivity. Intriguingly, even though variants with reduced biofilm formation ability displayed lessened root colonization in mono-culture, these were able to evolve and persist in most populations benefitting from the matrix overproduced by another morphotype. Finally, we observed increased root colonization of evolved isolates exclusively on *A. thaliana*. The lack of such increase on tomato (*Solanum lycopersicum*) roots suggested an *A. thaliana*-specific adaptation.

## Results

### B. subtilis shows rapid improvement and diversification during evolution on plant roots

To examine how the PGPR *B. subtilis* evolves on plant roots and adapts in root colonization, five parallel populations (from A to E), started from a single clonal lineage of *B. subtilis* strain DK1042 (natural competent derivative of the undomesticated NCBI 3610 (Konkol *et al*., 2013)), were passaged on *A. thaliana* seedlings for 20 transfers, spanning over a total of 40 days (Fig. 1). To follow possible improvements in root colonization during the experimental evolution, colony forming unit (CFU) per gram of roots was assessed after transfer 1, 5, 10, 17 and 20 (T1, T5, T10, T17 and T20, respectively). As expected, root colonization improved quickly in all evolving populations. The number of cells colonizing the roots increased on average about 10-fold, considering all five populations, from the first to the 5^th^ transfer and stayed at similar level afterwards (Fig. 2A). Tukey’
ss test confirmed that T5, T10 and T20 were significantly different from T1, while T17 was not significantly different from any of the others.

**Fig. 1.**
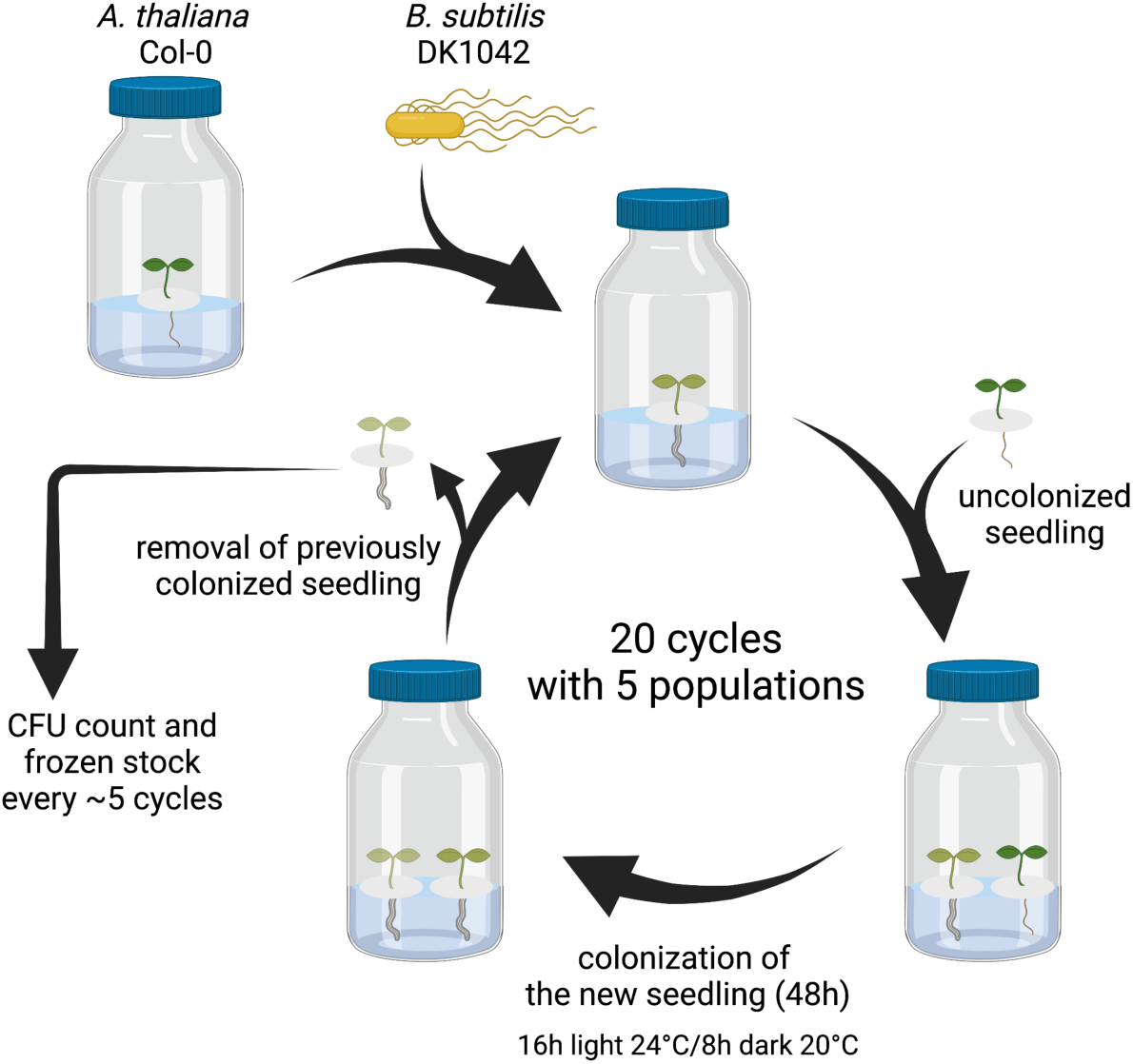
Experimental evolution scheme of parallel populations of *B. subtilis* (see Experimental procedures for detailed description of conditions used). Created with BioRender.com.

**Fig. 2.**
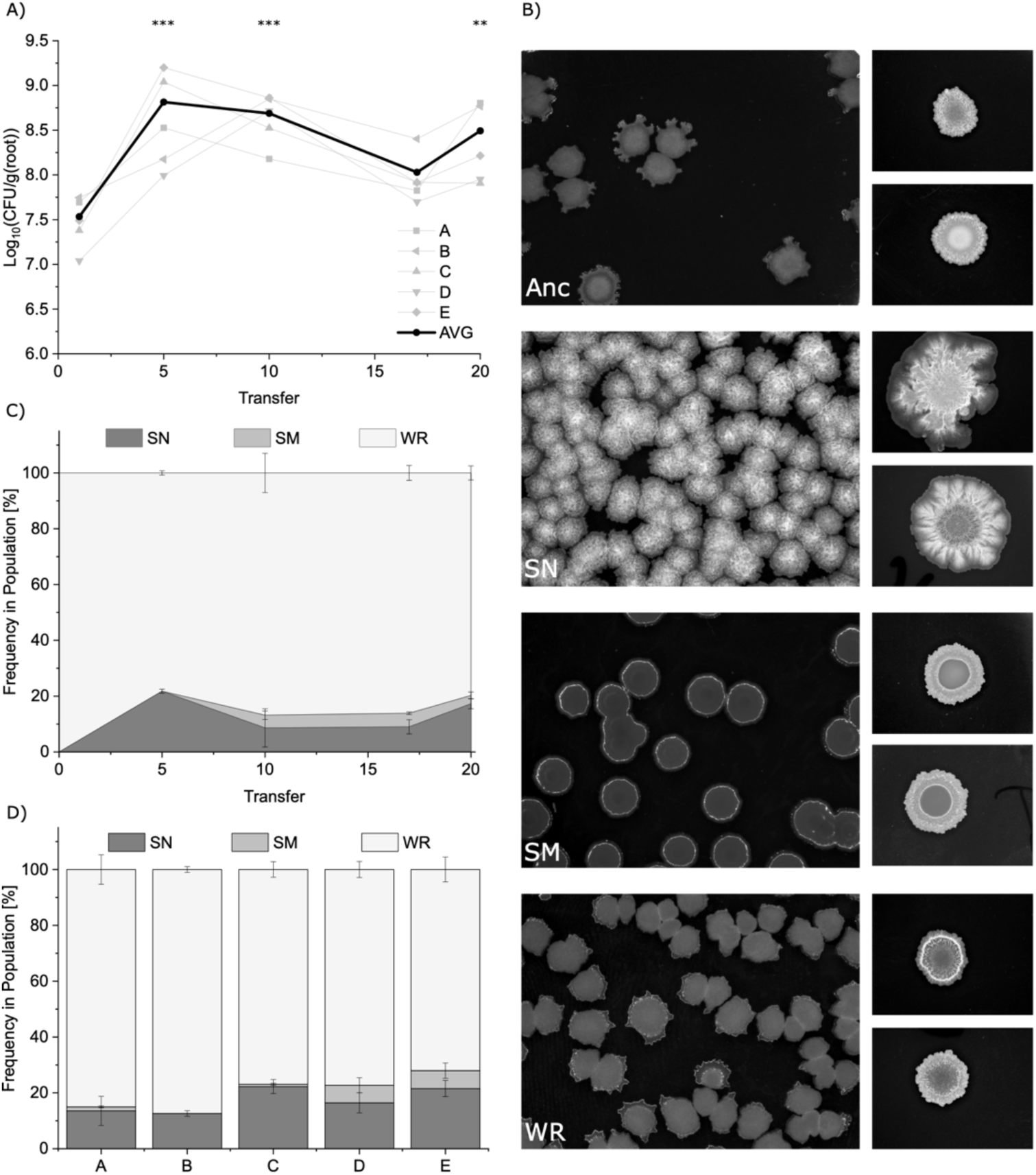
Adaptation in root colonization and diversification of *B. subtilis* on *A. thaliana*. A. Changes in root colonization during experimental evolution was monitored as CFU per gram root for T1, T5, T10, T17 and T20 in populations A to E. (**: p<0.01, ***: p<0.001). B. Distinct colony morphologies were identified and described based on colony architecture when plated (left) or spotted (right, two representative samples) on 1.5% LB-Agar after 48h incubation at 30°C for the ancestral strain (Anc), snow morphotype (SN), smooth morphotype (SM) and wrinkled morphotype (WR). C. Relative frequencies of different morphotypes in subsequent evolutionary time points in average of all five populations (n=5). D. Final (T20) relative frequencies of different morphotypes in all five parallel evolved populations (n=3). For panel C and D, data represent mean and error bars represent standard error.

Furthermore, in all five populations, three very distinct morphological variants were observed in the CFU assays. These include a Snow (SN) variant with hyper-wrinkled, whitish colony morphology, a Smooth (SM) variant with round, smooth and shiny colony morphology and a Wrinkled (WR) variant with a colony morphology identical to the ancestor (Fig. 2B). Because WR variants were indistinguishable from the ancestor by sheer colony morphology, only cultures directly from the ancestral stock were regarded and termed as ancestor in this report, while all other colonies and cultures, i.e. those that were observed after the first transfer of experimental evolution, were counted as WR. Importantly, identification of morphotypes via serial dilution plating on LB-agar has its limitations in the range of detection, therefore, the presence of morphotypes in low abundances under detectable levels cannot be excluded. Genetic stability of the three distinct morphotypes was confirmed by repeated passaging on either solid medium, in liquid culture or alternating between the two for a chosen representative example of each morphotype isolated from three populations (population A, D, and E). SN was observed in all five populations at T5 reaching in average a final frequency of around 20% (Fig. 2CD). SM was observed in all five populations at T10, except for population E, where it was observed after T15. The final frequencies of the morphotypes were highly similar across all five populations, except for population B, where SM was below detection level at T20 (Fig 2D).

### Pellicle formation and growth on plant polysaccharides correlate with distinct morphotypes

To further explore the phenotypic traits of the three distinct morphotypes, morphologically distinct isolates from the final transfer were randomly selected, and tested for growth and biofilm pellicle formation in response to plant polysaccharides. As *B. subtilis* was evolved in a minimal salt medium (see Experimental procedures), bacterial cells were dependent on carbon sources provided by the plant for their growth. We hypothesized, therefore, that evolved isolates could adapt to the environment by improving the utilization of plant polysaccharides. Indeed, most evolved isolates showed altered planktonic growth in MSNc + 0.5% xylan. Growth patterns correlated well with the specific morphotypes, i.e. prolonged stationary phase/delayed decline of SN, and higher growth rate as well as higher maximum optical density (OD) of SM compared to the ancestor (Fig. 3A). Notably, xylan displays some absorbance at OD_600_. Therefore, the initial drop in OD_600_ can be attributed to the time span where cells break down xylan, reducing OD_600_ until bacterial growth is able to exceed (Beauregard *et al*., 2013).

**Fig. 3.**
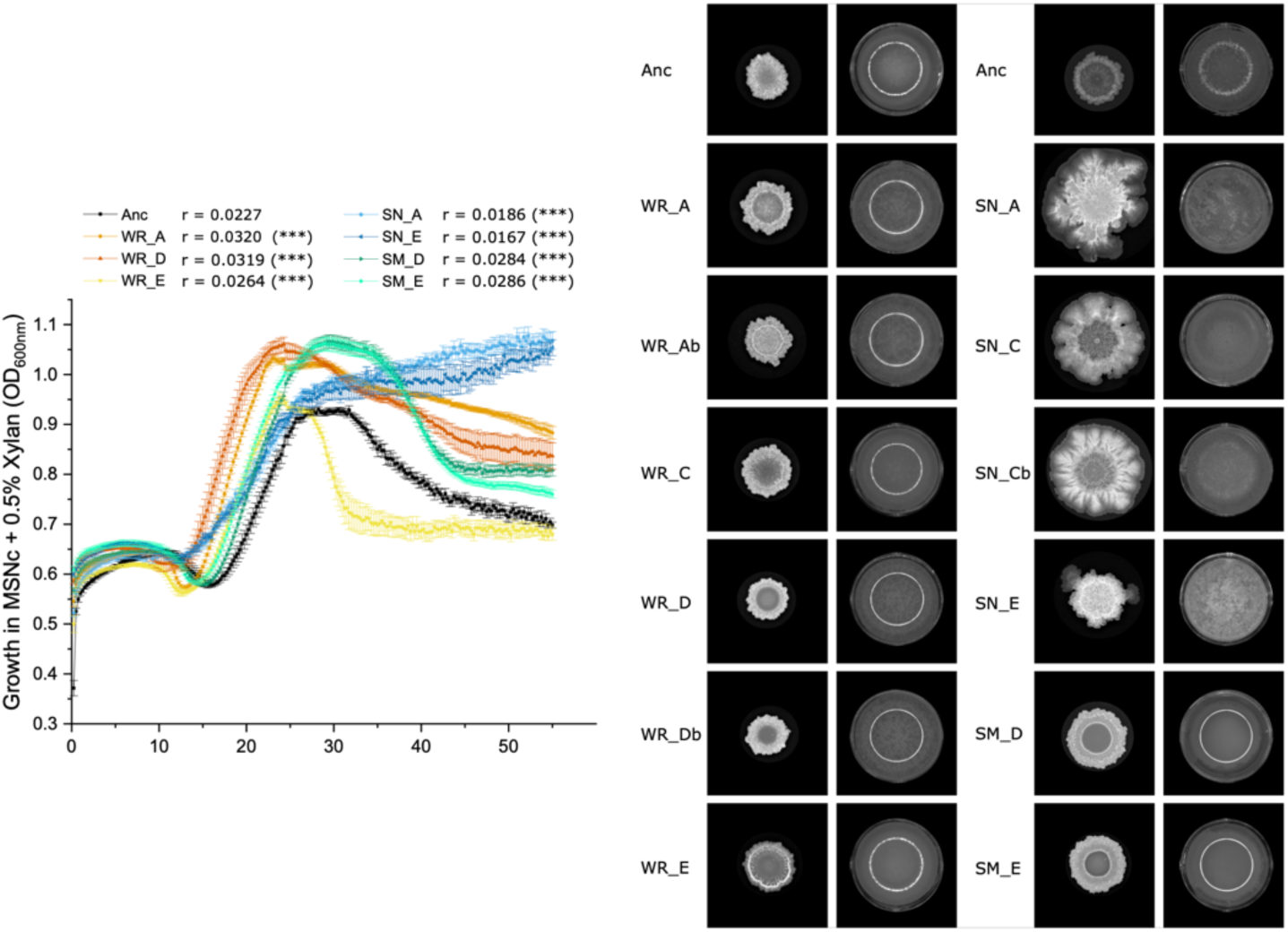
Growth and pellicle formation of evolved isolates from final transfer (T20) in response to plant polysaccharides compared to the ancestor. A. Growth of evolved isolates and ancestor in MSNc + 0.5% xylan was followed at 24°C for 56h (n=6, 2 independent overnight cultures with each 3 replicates). Data represent mean and error bars represent standard error. Growth rates (r) have been calculated for exponential phase following a linear regression model (***: p<0.001). B. Colony and pellicle morphology developed by evolved isolates and ancestor. Colony morphology was tested by spotting 2µl of ON culture on 1.5% LB-Agar and incubating for 48h at 30°C. Pellicle morphology was studied by inoculating overnight cultures into 1 mL LB medium supplemented with 0.5% xylan at a starting OD_600_ of 0.075. Images of colonies and pellicle were obtained after 48h at 30°C.

Pellicle formation has been proven to be not only a great indicator for biofilm productivity in general, but also to correlate with root colonization by *B. subtilis* strains (Chen *et al*., 2013). Beauregard *et al*. (2013) demonstrated that plant polysaccharides act as a signal for biofilm formation in *B. subtilis* via kinases of the Spo0A pathway. Xylan has been observed to be one of the strongest biofilm inducer plant polysaccharides (Beauregard *et al*., 2013). Here, we tested pellicle formation in LB medium in the presence of the plant polysaccharide xylan in 24-well plates. This allowed us to test the inductive effects on biofilm formation of *B. subtilis* by xylan, while still providing enough nutrients for robust pellicles. After incubating for 48h at 30°C, pellicle complexity and architecture of evolved isolates were analyzed in comparison to the ancestral strain. As expected, pellicle formation by the evolved isolates mostly correlated with morphological appearance of colonies on solidified LB medium (Fig. 3B). Similar to their colony morphology, SN variants produced hyper-robust pellicles, indicating overproduction of matrix components (Fig. 3B). On the contrary, SM variants were unable to form a stable pellicle, only a thin film was observed on the liquid-air interface (Fig. 3B). Notably, no major changes between pellicles formed by WR compared to the ancestral strain were observed (Fig. 3B).

### Improvements in plant root colonization tend to be A. thaliana specific

Biofilm formation on the root by *B. subtilis* is initiated by the recognition of plant signals, and the composition of those signaling components is suggested to be plant species-specific (Warembourg *et al*., 2003; Beauregard *et al*., 2013). Therefore, we wondered whether improvements in root colonization might be specific and limited to the plant species they evolved on. To test this, we inoculated both 14-16d old *A. thaliana* seedlings as well as 6-8d old tomato seedlings with either our evolved isolates from the final transfer or with the ancestral strain and quantified plant colonization after 48h.

Intriguingly, we observed that 7 out of 12 isolates showed improved root colonization compared to the ancestral strain on *A. thaliana*, while none improved on tomato (Fig. 4). This indicates that, *B. subtilis* adapted specifically to root colonization of *A. thaliana*. Interestingly, improvements in root colonization tended to be only partially in accordance with the morphotypes of the tested isolates. All four tested SN isolates, displayed improved root colonization on *A. thaliana*. This is consistent with previous studies, which showed that enhanced biofilm formation leads to increased root colonization (Chen *et al*., 2013). On the contrary, WR and SM isolates showed varying results. Remarkably, only SM_E showed reduced root colonization on both plant species in accordance with highly reduced pellicle formation, while SM_D performed similar to the ancestor. Notably, the two SM isolates evolved in distinct populations and might therefore harbor distinct mutations, therefore, additional mutations in the two SM isolates could explain variation in root colonization.

**Fig. 4.**
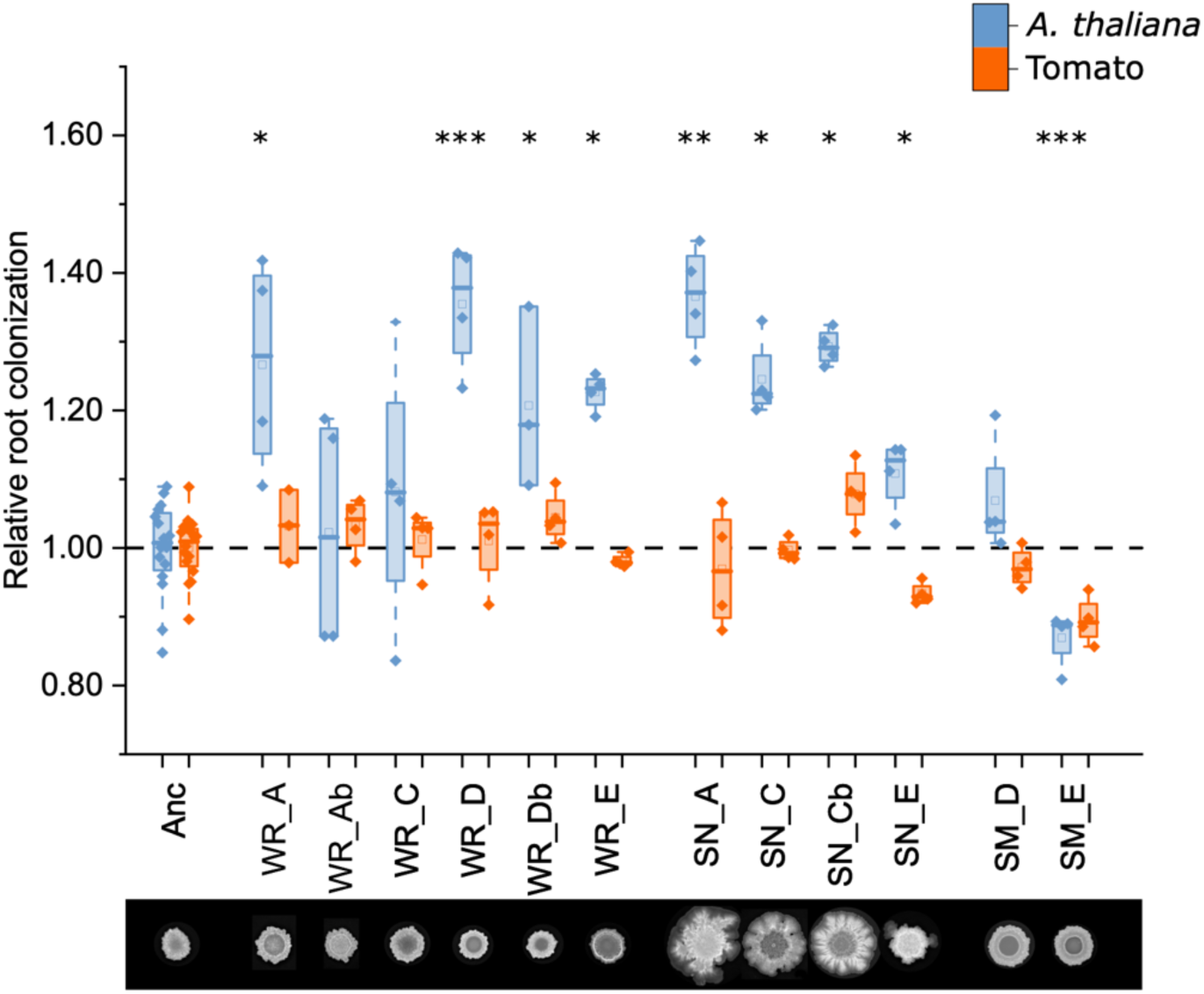
Root colonization by evolved isolates from the final transfer (T20) compared to the ancestor. Both 14-16 days old *A. thaliana* (blue) and 10-12 days old tomato seedlings (orange) were inoculated with evolved isolates originating from different populations (populations A, C, D, and E) as well as the ancestor and incubated for 48h in a plant chamber. The biofilm on the roots was disrupted through vortexing with small glass beads and the suspension was plated on LB-agar medium to allow CFU counting. For each isolate, relative root colonization was calculated by dividing the log10-transformed CFU value of each replicate with the mean of the log10-transformed CFU of the ancestor from the same experimental setup. Here, the pooled CFU of the ancestor from all experimental set-ups is shown. The dashed line represents the mean of the ancestor. ANOVA followed by Post-hoc Dunnett’
ss tests were performed to test for significance between each isolate and the ancestor (*: p<0.05, **: p<0.01, ***: p<0.001).

### Diversification involves mutations related to biofilm development, motility and metabolic pathways

To get an idea of the genetic basis behind the observed diversification, genomes of selected SN (SN_A, SN_E), SM (SM_D, SM_E) and WR (WR_A, WR_D and WR_E) isolates were re-sequenced. The genome of the ancestor was previously re-sequenced (Dragoš *et al*., 2018b), that revealed differences in the genome sequence of our laboratory DK1042 stock compared to the database. In total, 43 non-overlapping mutations (SNPs) were detected in the seven re-sequenced genomes, of which 38 were single-nucleotide variants, 3 were deletions, and 2 were insertions (Table 1). Both SN_A and SN_E harbored a single mutation, located in the gene coding for the biofilm master regulator, SinR. SM_D and SM_E shared a mutation in the *flhB* gene, important for flagellar assembly, as well as numerous mutations in the *gtaB* and *yvzE* genes, coding for UTP-glucose-1-phosphate uridylyltransferases. Each of them harbored additional SNPs, a non-synonymous mutation in the glycerol facilitator coding gene, *glpF*. Further, a synonymous mutation in *ydaD* gene was detected in SM_D, while SM_E additionally carried non-synonymous mutations in the sporulation specific genes, *spoVR* and *dacB*. Finally, the distinct WR isolates from different evolved populations shared no overlapping mutations, but commonly harbored SNPs in genes related to motility (*fliI, fliR*, and *motB* in WR isolates from population A, D, and E, respectively). WR_A and WR_D contained mutations in *pdeH* encoding a c-di-GMP degrading phosphodiesterase, next to few additional mutations in these isolates. Because SM isolates, evolved in independent populations, shared multiple identical mutations, genetic heterogeneity might have already be present in the ancestral starting culture. Nonetheless, the rise to detectable ratios of SM during the evolution indicates that they are advantageous for the population and are, thus, selected for over time.

**Table 1.** List of mutations detected in the genomes of re-sequenced morphotypes. Functions of the gene products were retrieved from SubtiWiki Database (Zhu & Stülke, 2017)

| SN |  | SM |  | WR |  |  | Position | Gene | Mutation | Function of gene product |
| --- | --- | --- | --- | --- | --- | --- | --- | --- | --- | --- |
| SN_A | SN_E | SM_D | SM_E | WR_A | WR_D | WR_E |  |  |  |  |
|  |  | ✓ |  |  |  |  | 472036 | *ydaD* | 288G>T | general stress protein |
|  |  |  |  | ✓ |  |  | 478790 | *intergenic* | C>G | upstream region of *ydaJ* that codes for a lipoprotein that may modify the extracellular polysaccharide synthesized by YdaLMN proteins |
|  |  |  |  |  | ✓ |  | 478876 | *intergenic* | T>del |  |
|  |  | ✓ |  |  |  |  | 1002770 | *glpF* | Pro78Ala | glycerol facilitator |
|  |  |  | ✓ |  |  |  | 1016370 | *spoVR* | Glu229Gly | spore cortex synthesis |
|  |  |  |  | ✓ |  |  | 1391100 | *thiU* | Pro192Gln | thiamine uptake |
|  |  |  |  |  |  | ✓ | 1433817 | *motB* | Gly226Asp | flagellar motor rotation |
|  |  |  |  | ✓ |  |  | 1696818 | *fliI* | Thr304Lys | flagellar-specific ATPase |
|  |  |  |  |  | ✓ |  | 1706108 | *fliR* | Ala89_Ala93del | part of the flagellar Type III secretion system |
|  |  | ✓ | ✓ |  |  |  | 1707666 | *flhB* | Glu347fs | part of the flagellar Type III secretion system |
|  |  |  | ✓ |  |  |  | 2424537 | *dacB* | Ala7Thr | sporulation-specific penicillin-binding protein |
| ✓ | ✓ |  |  |  |  |  | 2552926 | *sinR* | Ala85Thr | control of biofilm formation |
|  |  |  |  | ✓ |  |  | 2578188 | *pstBA* | Ser216Pro | phosphate ABC transporter ATP-binding protein |
|  |  |  |  |  | ✓ |  | 3258864 | *pdeH* | Asp137fs | degradation of c-di-GMP |
|  |  |  |  | ✓ |  |  | 3258936 |  | Asp116Tyr |  |
|  |  | ✓ | ✓ |  |  |  | 3666149 | *gtaB* | 483G>A | UTP-glucose-1-phosphate uridylyltransferase |
|  |  | ✓ | ✓ |  |  |  | 3666155 |  | 489G>A |  |
|  |  | ✓ | ✓ |  |  |  | 3666161 |  | Glu165Asp |  |
|  |  | ✓ | ✓ |  |  |  | 3666171 |  | Arg169Cys |  |
|  |  | ✓ | ✓ |  |  |  | 3666173 |  | 507C>T |  |
|  |  | ✓ | ✓ |  |  |  | 3666176 |  | 510C>T |  |
|  |  |  | ✓ |  |  |  | 3666182 |  | 516T>C |  |
|  |  | ✓ | ✓ |  |  |  | 3673066 | *yvzE* | Phe27Val | putative UTP-glucose-1-phosphate uridylyltransferase |
|  |  | ✓ | ✓ |  |  |  | 3673068 |  | Phe27Leu |  |
|  |  | ✓ | ✓ |  |  |  | 3673077 |  | 90A>T |  |
|  |  | ✓ | ✓ |  |  |  | 3673098 |  | Asp37Glu |  |
|  |  | ✓ | ✓ |  |  |  | 3673101 |  | 114A>T |  |

### Complementarity leads to enhanced productivity of mixed populations in root colonization

Finally, we wondered how the SM isolates, which showed reduced pellicle biofilm development in general and significantly reduced root colonization for one isolate, rose to and maintained low frequencies throughout the evolution experiment and was not outcompeted over time during the evolution. We hypothesized that its root colonization in the evolving populations might be facilitated by other co-occurring morphotypes. Therefore, we tested the impact of diversity on individual and group performance in root colonization by inoculating *A. thaliana* seedlings with SN, SM and WR in monoculture as well as with a mix of all three morphotypes with equal starting frequencies (1:1:1) for three different populations each (populations A, D, and E). We used an established method to distinguish between selection, where variants that are most productive in monoculture tend to dominate the mix, and complementarity, where most or all constituents tend to benefit from biodiversity (Loreau & Hector, 2001; Poltak & Cooper, 2011).

Even though the mix of all three morphotypes tended to perform better in all three populations than the ancestor (p_PopA_ = 0.0220, p_PopD_ = 0.0620, p_PopE_ = 0.0055), its CFU was the overall highest only in population D, while in population A and E, it was lower than SN in monoculture, even though not significantly (Fig. 5). However, the observed productivity of the three mixes were higher than their respective predictions. To this end, a complementarity effect of biodiversity seemed to be the driving force affecting the productivity of mixed populations. In population D, all three morphotypes appeared to benefit from each other, as the CFU of each of the morphotypes were higher in the co-culture than predicted, suggesting a mutualistic interaction. Productivity of SN increased thereby by 1.96-fold, SM by 2.50-fold and WR by 4.50-fold relative to their predicted CFU in the mix, resulting in 103% complementarity and -3% selection. On the contrary, in population A and E only SM and WR benefited from the co-culture, while SN showed highly reduced CFU in the mix compared to its prediction. In populations A and E, productivity of SN decreased by 0.18-fold and 0.47-fold, while it increased for SM by 2.50-fold and 1.70-fold and WR by 20.72-fold and 4.75-fold, leading to 641% and 375% complementarity as well as -541% and -275% selection, respectively.

**Fig. 5.**
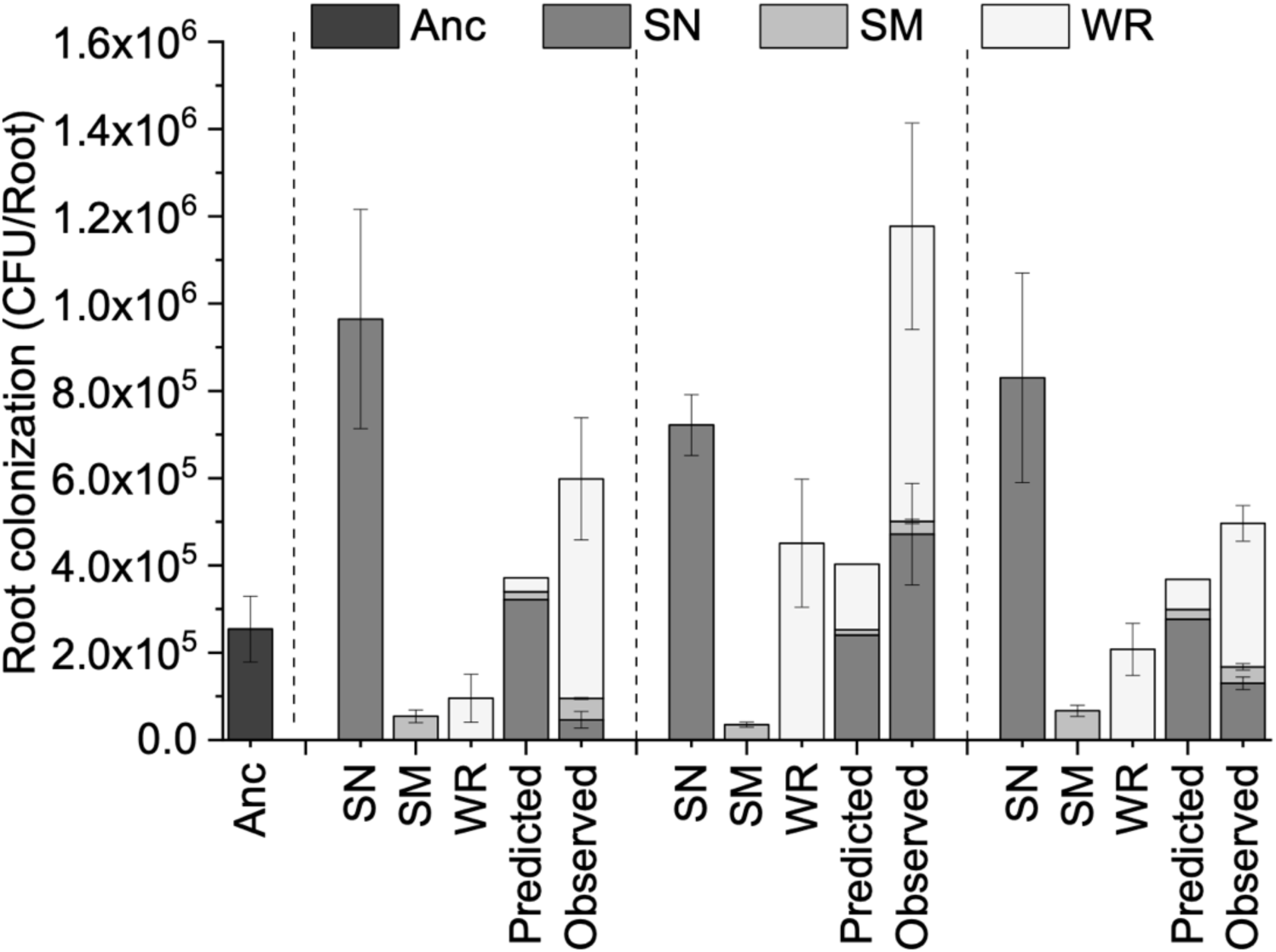
Root colonization of morphotypes in monoculture as well as in a 1:1:1 mix or the ancestor. 14-16d old *A. thaliana* seedlings were inoculated with either the mix or the strains in monoculture and incubated for 48h in a plant chamber. The biofilm on the roots was disrupted by vortexing with small glass beads and the suspension was plated on solidified LB medium to allow CFU counting. Predicted colonization was calculated as the product of the proportion of each morphotype in the inoculum (1/3) and its performance (CFU per root) in monoculture. Observed colonization is the total colonization of the mixed community in the experimental setup. Bars represent the mean and error bars represent standard error for four replicates (9 replicates for ancestor pooled from different days).

## Discussion

The PGPR *B. subtilis* is a promising sustainable alternative to traditionally used agrochemicals, as it harbors many important plant growth-promoting and biocontrol properties while having reduced negative effects on biodiversity and the environment (Blake *et al*., 2021; Arnaouteli *et al*., 2021). Even though its potential seems immense under laboratory conditions and in greenhouse trials, the success varies greatly when applied in fields (Peng *et al*., 2011; Bardin *et al*., 2015; Moreira & de Mio, 2015; Wei *et al*., 2016). To improve the application of *B. subtilis* within agriculture, we need a deeper understanding of the underlying mechanisms of how it interacts with and colonizes plants. In this work, we used experimental evolution to analyze how *B. subtilis* evolves on *A. thaliana* roots, by continuously transferring a previously colonized root to a 100ml reagent bottle containing fresh medium and a sterile root, thereby allowing only root-colonizing cells to be carried on throughout the experiment. In contrast to the experimental evolution study by Batstone *et al*. (2020) that similarly examined plant-microbe interaction, we focused solely on initial and re-colonization of plant roots by *B. subtilis* in liquid minimal media with small plant seedlings. By using MSNg, a minimal media with low glycerol concentrations that is insufficient to support growth and biofilm development of a *B. subtilis* monoculture (Beauregard *et al*., 2013), we promote bacteria being dependent on carbon provided by the plant and possibly also being stimulated to move towards the root. Thus, by passaging only the seedlings, and therefore only cells that attach tightly to the roots, we were able to select for root colonization. Because traits such as motility, biofilm formation and dispersal as well as the recognition and utilization of plant polysaccharides are assumed to be important and beneficial for root (re-)colonization in our experimental set-up, we expected cells to improve in these properties over time (Chen *et al*., 2013; Allard-Massicotte *et al*., 2016; Dragoš *et al*., 2018a). Indeed, we observed that *B. subtilis* populations rapidly increased in root colonization during the experimental evolution, indicating fast adaptation to plant roots. In order to test whether genetic adaptation had taken place, single evolved isolates from the final transfer (T20) were randomly selected for further characterization, including growth and biofilm formation in response to plant polysaccharides. Intriguingly, when testing evolved isolates from the final transfer for improved root colonization of *A. thaliana* and tomato seedlings, we only observed improvements on *A. thaliana* indicating *A. thaliana*-specific adaptation paths. Furthermore, we observed that evolving populations quickly diversified into three distinct morphotypes, comprising a hyper-wrinkled SN variant, a round and smooth SM variant, and a variant identical in colony morphology to the ancestor, termed WR. Finally, mixed colonization of distinct morphotypes revealed a complementarity effect of biodiversity during root colonization, as the performance of the mix exceeded the prediction based on individual performance of its constituents (Fig. 5). Depending on the evolved population they originated from, SN, SM and WR interacted either in a mutualistic way, as all three morphotypes benefited, or SM and WR most likely exploited SN by capitalizing on overproduced matrix components.

Previous work proved that rapid diversification is commonly observed in microbial evolution and also occurs in experimental evolution of *B. subtilis* in liquid LB culture under static and shaken conditions as well as in biofilms (Leiman *et al*., 2014; Dragoš *et al*., 2018b; Richter *et al*., 2018). Here, we demonstrated that diversification of *B. subtilis* also plays an important role in experimental evolution during ecologically- and biotechnologically-relevant plant root colonization. Diversification of *B. subtilis* during evolution on *A. thaliana* roots seems highly reproducible, as the same evolutionary pattern and highly similar morphotype frequencies could be observed in the final transfer of all five parallel lines. The most striking difference between distinct morphotypes occurred in the complexity in colony and pellicle architecture, indicating alterations in the capability of biofilm formation. In addition to the hyper-wrinkled SN variant, from which all tested isolates showed hyper-robust biofilm formation in the form of complex colonies and thick pellicles, a SM morphotype emerged that seemingly had lost its ability to form a stable pellicle when inoculated solitarily. In accordance with previous work by Chen *et al*. (2013), who tested correlation of pellicle formation with root colonization of distinct natural isolates of *B. subtilis*, the evolved isolates displaying a more robust pellicle formation, namely SN, also improved in root colonization. On the contrary, only one of the tested SM isolates showed reduced root colonization, while unexpectedly the other SM isolate performed similar to the ancestor on plant roots. Notably, dissimilar traits of the analogous morphotypes could be either caused by additional mutations or alternatively, distinct mutations led to similar morphological, but slightly different plant colonization consequences (Poltak & Cooper, 2011).

Remarkably, evolved isolates showed improvements in root colonization only on *A. thaliana*, but not on tomato. In nature, the plant host species tend to be important drivers for diversity of the plant microbiome, indicating active selection by the plant (Compant *et al*., 2019). Furthermore, PGPR have been shown to respond species-specifically in chemotaxis towards their original host plant where these were isolated from (Zhang *et al*., 2014). Distinct plant species produce and secrete distinct arrays of root exudates, which act not only as a major carbon source for PGPR, but also as chemical signals for bacterial chemotaxis and are, thus, important initiators of root colonization (Warembourg *et al*., 2003; Beauregard, 2015). Similar to root exudates, the composition of plant polysaccharides, which are known inducers of biofilm formation, differs between plant species (Beauregard *et al*., 2013). Lastly, the sheer differences in the physiological architecture of plant roots, including size, thickness, abundance of lateral roots and root hairs, might provide even more variation in environmental conditions between distinct plant species, thereby allowing plant species-specific adaptations (Saleem *et al*., 2018). Although the Arabidopsis-specific aspects of the adaptation mechanism still need to be revealed, the evolved isolates performed differently on the two plant species.

Even though this report focuses mainly on phenotypical aspects of evolution of *B. subtilis* on *A. thaliana* roots, we aimed to get a first impression of the genetic basis behind the adaptive mechanisms, and consequently re-sequenced genomes of seven evolved isolates. In both of the tested SN isolates, a mutation in the gene encoding the biofilm regulator SinR (A85T) was found. An identical mutation in the *sinR* gene was previously observed resulting in hyper-wrinkled colony morphology, increased expression of biofilm-related genes and enhanced matrix production (Chai *et al*., 2010; Leiman et al., 2014; Richter *et al*., 2018; Kampf *et al*., 2018). Intriguingly, both Richter *et al*. (2018) and Kampf *et al*. (2018), observed spontaneous evolution of the exact same point mutation A85T in *sinR*, indicating strong positive selection. Further, *ΔsinR* mutant *B. subtilis* with hyper-robust biofilm displayed increased number of cells attached to plant roots (Chen *et al*., 2013). These previously reported data indicate that the observed mutation in *sinR* results in enhanced matrix production, biofilm formation. Ultimately, enhanced biofilm matrix production resulted in increased root colonization of the SN isolates. Unfortunately, observed mutations in SM and WR isolates are not as self-explanatory as in SN, and, thus, we can only speculate about their consequences requiring further molecular validation. The sequenced SM isolates harbored multiple mutations in *gtaB* and *yvzE* genes, coding for UTP-glucose-1-phosphate uridylyltransferases synthesizing a precursor for exopolysaccharide biosynthesis (Varon *et al*., 1993). Intriguingly, acetylation of GtaB was demonstrated to be important for proper biofilm formation in *B. subtilis* and a *gtaB* deletion mutant showed reduced pellicle development (Reverdy *et al*., 2018), suggesting that *gtaB* mutations could be relevant in the observed SM morphotypes. In addition, we speculate that the multiple mutations in *gtaB* possibly resulting in an unfunctional protein could explain the increased growth of the SM isolates due to decreased EPS production. Two of the three WR variants carried distinct mutations in the *pdeH* gene that encodes a c-di-GMP degrading phosphodiesterase, and additional, again distinct, mutations in the intergenic region upstream of *ydaJ* that encodes a lipoprotein that may modify the extracellular polysaccharide synthesized by YdaLMN proteins (Bedrunka & Graumann 2017). Notably, perturbation of these genes slightly alters colony morphologies (Bedrunka & Graumann 2017). Finally, all SM and WR morphotypes carried mutations in genes related to motility. While motility is not essential for biofilm formation, motile strains are benefited in biofilm development compared to non-motile derivates (Hölscher *et al*., 2015), therefore these mutations might additionally impact the efficiency of plant colonization. Nevertheless, overexpression of the *eps* operon in the SN morphotypes due to *sinR* mutation, could also possibly impact motility due to the EpsE clutch (Blair *et al*., 2008). However, the role and impact of motility on the evolution of *B. subtilis* during plant root colonization will be examined in future studies.

The observations and results from monoculture experiments are not able to explain how and why SM variants, diminished in their ability to form biofilms and partially reduced in root colonization, are able to arise and persist in populations during the evolution. Hence, we expected SM to capitalize on the presence of other co-occurrent morphotypes. A general observation from several experimental evolution studies with biofilm models is that the productivity of evolved populations not necessarily equals to the sum of productivities of its members in monocultures, as interactions between distinct evolved variants might occur (Brockhurst *et al*., 2006; Poltak & Cooper, 2011; Ellis *et al*., 2015). Overall, biodiversity has been regarded to impact productivity of a community, by two distinct processes, namely selection and complementarity (Loreau & Hector, 2001). Selective processes are anticipated to lead to dominance of community members, which perform best in monoculture, as they are expected to be able to outcompete others (Loreau & Hector, 2001). Complementary processes comprise advantageous effects of biodiversity, where most or all constituents benefit (Loreau & Hector, 2001). Interactions in a mixed community might, therefore, not always be positive, but can be also negative, as either mutualistic interactions, e.g. cross-feeding and niche construction, or antagonistic interactions, e.g. cheats and exploitation, might evolve (Day & Young, 2004; Poltak & Cooper, 2011, Dragoš *et al*., 2018b). Here, we extend these observations with a biotechnological relevant system on plant roots, describing the evolution of both exploitative as well as mutualistic interactions between the evolved morphotypes in distinct evolutionary lineages. The number of cells colonizing plant roots with an initially equal mix of SN, SM and WR is higher compared to its prediction in all three tested evolutionary lineages. Indeed, a complementarity effect of biodiversity seemed to be the cause for this “overyielding” of the mixed community. However, only in one of the three tested populations, namely in the mix of population D, all three morphotypes seemed to benefit from being in this mix, indicating mutualistic interactions. Further, this was the only tested population, were the mix outperformed all monocultures. In the other two tested evolved populations, even though the mix performed better than the ancestor, the three members population did not outperform the best performing monoculture, namely SN. Additionally, SN showed strongly reduced CFU in these two mixes compared to its predicted value, suggesting that only WR and SM benefited. The observed mutation in *sinR* in SN suggests that it overproduces matrix components, a public good that is costly, but can be capitalized on. We speculate that WR and SM possibly exploit this, by integrating into the matrix without having the cost of its production, improving their own productivity in mixed colonization at the expense of SN. Considering that SM was observed during experimental evolution only after SN rose to detectable levels in each of the five parallel populations, it is reasonable to infer that SM is able to arise and persist in evolved populations by capitalizing on the enhanced matrix provided by SN. Similar to our study, Dragoš *et al*. (2018b) observed the evolution of a non-producing and matrix-exploiting smooth morphotype in *B. subtilis* pellicles. The described smooth variant integrated and resided in the biofilm produced by co-occurring morphotypes, thus, exploiting matrix production (Dragoš *et al*., 2018b).

Even more significantly, complementary effects are observed during root colonization by distinct morphotypes, evolved from a shared ancestor, in all tested populations. Indeed, a negative selection effect of biodiversity, as observed for all three populations here, indicates that the best root colonizing variant in monoculture is losing out in a mixed community. This requires attention in future studies, as it raises many questions about the influence of the natural diversity in the rhizosphere on the application of *B. subtilis*. Further, the observation that interactions between the three morphotypes differed in distinct evolved populations indicates that biodiversity both evolved and is maintained differently in the parallel populations (Poltak & Cooper, 2011). Nevertheless, further experiments are needed to reveal whether similar ecological interactions exist among the isolates from distinct lineages.

In conclusion, our results suggest that *B. subtilis* adapts rapidly to root colonization on *A. thaliana*. Our findings are relevant for the application of *B. subtilis* as a PGPR in agriculture, as they imply that genetic diversification might be a stable strategy for root colonization, and a mix of distinct phenotypic variants might be more beneficial compared to one single strong root colonizer. In addition to a potential increase in net productivity on plant roots, a diverse mix might be more resistant to invasions by non-cooperating cheaters (Brockhurst *et al*., 2006), therefore, enhancing also the maintenance of the biofilm formed on the root.

## Experimental procedures

### Bacterial strains and culture conditions

Throughout this study, the ancestral strain *B. subtilis* DK1042 (natural competent variant of the undomesticated NCIB 3610 (Konkol *et al*., 2013)) and evolved derivatives were grown in Lysogeny Broth (LB) (Lennox, 10 g·l^-1^ Tryptone, 5 g·l^-1^ Yeast Extract and 5 g·l^-1^ NaCl) over night (ON) at 37°C. For formation of complex colonies, 2µl of exponential phase culture were spotted on LB medium solidified with 1.5% agar and incubated at 30°C for 48h. To assay pellicle formation, the OD_600_ of a bacterial culture in exponential phase was adjusted to 5, and 15µl were inoculated into 1ml of LB + 0.5% xylan (from beechwood, Carl Roth) in 24-well microtiter plates and incubated at 30°C for 48h. In order to analyze growth in response to plant polysaccharides, the OD_600_ of a bacterial culture in exponential phase was adjusted to 0.1 in a minimal salt medium (MSN) (5mM potassium phosphate buffer pH 7.0, 0.1M 3-(N-morpholino)propanesulfonic acid pH 7.0, 4 mmol·l^-1^ MgCl_2_, 0.05 mmol·l^-1^ MnCl_2_, 1 μmol·l^-1^ ZnCl_2_, 2 μmol·l^-1^ thiamine, 700 μmol·l^-1^ CaCl_2_, 0.2 % NH_4_Cl) supplemented with 0.5% cellobiose (Carl Roth) (MSNc) and 0.5% xylan, because xylan and cellulose have been previously observed to support growth of *B. subtilis* (Beauregard *et al*., 2013). 100µl was transferred to a 96-well plate and incubated in a plate reader (BioTek Synergy HTX Multi-Mode Microplate Reader) for 58h at 24°C, orbital shaking. OD_600_ was measured every 15 minutes in a 9-point measurement at different points of each well, to exclude errors of clumping and inhomogeneous suspensions. Afterwards, each well was analyzed under the stereomicroscope to confirm no visible aggregation or pellicles formed, which would skew OD_600_ measurements. Exponential phase of growth curves was identified with a linear regression model and growth rates (r) were calculated. For root colonization experiments MSN supplemented with 0.05% glycerol (MSNg) was used.

### Seedling preparation

In this study, both *A. thaliana* ecotype Col-0 and tomato (*Solanum lycopersicum L*., Maja Buschtomato, Buzzy Seeds, NL) seedlings were used. Seeds were surface sterilized in Eppendorf tubes by shaking vigorously with an orbital mixer for 12 minutes in 1mL 2% sodium hypochlorite. Afterwards, seeds were washed five times in 1 ml sterile MiliQ water through alternating centrifugation and removal of liquid solution. After sterilization, 8-10 seeds were plated on pre-dried 1% MS-agar (Murashige and Skoog basal salts mixture; Sigma) (2.2 g l^−1^) ensuring that the seedlings have sufficient space for proper root development and plates were sealed with parafilm. *A. thaliana* seeds were stratified in the dark at 4°C for 2-3d and subsequently incubated in a plant chamber with a day/night cycle of 16h light at 24°C and 8h dark at 20°C for 12-14d at an angle of 65°. Tomato seeds were moved to the plant chamber without stratification and incubated for 6-8d.

### Experimental evolution

We established an experimental evolution approach inspired by the bead transfer model of Poltak & Cooper (2011) to study *B. subtilis* evolution on plant roots, similar to the setup exploited for *Bacillus thuringiensis* (Lin *et al*., 2020). Therefore, every other day, a previously colonized plant seedling was transferred to a new sterile seedling, allowing only bacterial cells that attach to the new root to continue in the experiment, enabling selection for a continuous cycle of dispersal, chemotaxis towards the plant root, and biofilm formation on the root. To start the experimental evolution, five 14-16d old *A. thaliana* Col-0 seedlings (population A to E) were placed in 100mL reagent bottles containing 27ml of MSNg, and inoculated with 3ml of bacterial culture of *B. subtilis* DK1042 adjusted to OD_600_ of 0.2, reaching a starting OD_600_ of 0.02. To ensure that only the roots but not sprout or leaves provided surface for colonization, the seedlings were pulled through a sterile nylon mesh floating on top of the liquid medium, leaving only the roots submerged in the medium (Harris *et al*., 2019). Seedlings were incubated for 48 hours under static conditions in a plant chamber with a day/night cycle of 16h light at 24°C and 8h dark at 20°C. Before the transfer, the mesh of the previously colonized root was removed and the seedling was carefully washed in sterile MilliQ water, in order to remove non-colonizing cells. Subsequently, the colonized seedling was placed in fresh medium containing a new sterile seedling, allowing re-colonization. Evolving populations were continuously passaged every 48h for a total of 40 days (20 subsequently colonized seedlings per population). In order to keep track and ensure sterile handling of seedlings during the evolution, two controls without initial inoculant were included, in which either sterile seedlings or 10% of the MSNg medium were passaged. To follow the development of the populations and test for contamination, the bacterial cells colonizing the old roots, as well as the two controls, were plated on LB agar medium after initial colonization and transfer 5, 10, 17 and 20. In order to disperse the biofilm from the seedling for the quantification, roots were detached from the sprout, carefully washed in sterile MilliQ, and vortexed vigorously for 3 minutes in 1ml of 0.9% NaCl with small glass beads (Glass beads Ø0.75-1.00 mm, Carl Roth). The resulting cell suspensions were plated for CFU quantification and saved as frozen stocks for later comparison.

### Morphological diversification

Distinct morphotypes of evolved populations were determined according to characteristics in colony morphology (shape, size and color) on LB agar medium. To ensure different morphotypes were genetically stable, three random colonies of each variant were isolated and passaged every 24h for three transfers either in liquid LB, on LB agar medium as well as alternating between the two media for four transfers. After the final transfer, cultures were serial diluted and plated on LB-Agar to assess colony morphology. In order to analyze the development of the ratio of each morphotype during the evolution, a bacterial culture prepared from the frozen stock of each population for transfer 5, 10, 17, and 20, as well as of the ancestor as starting point 0, was grown to log-phase, diluted, plated on LB-agar in triplicates for two dilutions and incubated for 16h at 30°C. Subsequently, morphotypes were identified, all colonies were counted, with a minimum of 50 and an average of 100 colonies per plate, and ratios were calculated (Leiman *et al*., 2014).

### Root colonization productivity

In order to analyze whether evolved isolates were improved in root colonization compared to the ancestor, and whether those improvements were specific for the plant species the isolates evolved on, root colonization was tested on both *A. thaliana* Col-0 and tomato seedlings. Therefore, three single isolates were picked randomly from the final populations (i.e. T20) of linages A, C, D, and E. 14-16d old *A. thaliana* and 6-8d old tomato seedlings were inoculated with each of the evolved isolates in four replicates as described previously for initiation of the experimental evolution. After confirmation of equal variance with Levene’
ss test and normality of data with Shapiro–Wilk test, changes in root colonization were analyzed for significance with Dunnett’s post hoc test. Furthermore, to test the impact of diversity on the individual and mixed performance in root colonization, 14-16d old *A. thaliana* seedlings were inoculated with a 1:1:1 mix of pre-grown cultures of the three distinct morphotypes (SN, SM and WR), originating from the same evolved population (population A, D and E). Random colonies of each of the three morphotypes have been newly isolated for population A and D, because one of the morphotypes lacked in either population. Further, this ensured avoidance of a potential bias from previous monoculture experiments caused by different performances of the two tested WR isolates, e.g. WR_A and WR_Ab, in each of the populations. For population E, previously tested SN_E, SM_E and WR_E were used. Seedlings were incubated together with the bacterial inoculant for 48h in the plant chamber to allow root colonization. Afterwards, seedlings were carefully washed in sterile MilliQ water, the biofilm formed on plant roots was disrupted and the bacterial suspensions were plated on LB agar medium to allow CFU quantification. For mixed colonization, the predicted root colonization was calculated by multiplying the root colonization of each constituent in mono culture with the ratio in the inoculum (1/3) (Loreau & Hector, 2001; Poltak & Cooper, 2011). Further, selection and complementarity effect have been calculated as described by Loreau & Hector (2001).

### Microscopy

Bright field images of whole pellicles and colonies were obtained with an Axio Zoom V16 stereomicroscope (Carl Zeiss, Germany) equipped with a Zeiss CL 9000 LED light source and an AxioCam MRm monochrome camera (Carl Zeiss, Germany).

### Statistical analysis

At least three replicates were conducted for all experiments. Statistical analysis was performed in R 4.0.3. In detail, Shapiro-Wilk test and Levene’s test were performed to test for normality and equality of variance, respectively. To test for outlier, Grubb’
ss tests were conducted. For comparison between ancestor and evolved isolates, as well as between the first and following transfers during experimental evolution, ordinary one-way ANOVA analysis corrected with Benjamini-Hochberg procedure and Dunnett’s tests were employed. In order to test for significance in differences between predicted and observed CFU on roots in mixed colonization, Student’s t-tests were performed.

### Genome resequencing and genome analysis

Resequencing of evolved isolates was performed as previously (Dragoš *et al*., 2018b; Dragoš *et al*., 2018c; Martin *et al*., 2020; Thérien *et al*., 2020; Gallegos-Monterrosa *et al*., 2021). Genomic DNA of selected evolved isolates from the final transfer was extracted from 2ml ON cultures using the EURx Bacterial and Yeast Genomic DNA Kit. Paired-end libraries were prepared using the NEBNext® Ultra™ II DNA Library Prep Kit for Illumina. Paired-end fragment reads were generated on an Illumina NextSeq sequencer using TG NextSeq® 500/550 High Output Kit v2 (300 cycles). Primary data analysis (base-calling) was carried out with “bcl2fastq” software (v2.17.1.14, Illumina). All further analysis steps were done in CLC Genomics Workbench Tool 9.5.1. Reads were quality-trimmed using an error probability of 0.05 (Q13) as the threshold. In addition, the first ten bases of each read were removed. Reads that displayed ≥80% similarity to the reference over ≥80% of their read lengths were used in the mapping. Non-specific reads were randomly placed to one of their possible genomic locations. Quality-based SNP and small In/Del variant calling was carried out requiring ≥8 × read coverage with ≥25% variant frequency. Only variants supported by good quality bases (Q ≥ 20) were considered and only when they were supported by evidence from both DNA strands in comparison to the *B. subtilis* NCIB 3610 genome and pBS plasmid (GenBank accession no. NZ_CP020102 and NZ_CP020103, respectively). Identified mutations in each strain are listed in Table 1. Raw sequencing data has been deposited to the NCBI Sequence Read Archive (SRA) database under BioProject accession number: PRJNA705297

## Conflict of interests

The authors declare that there is no conflict of interests in relation to the work described.

## Notes

### Competing Interest Statement

The authors have declared no competing interest.

### Summary of Updates

new figure added and significance now indicated for certain data; manuscript updated

